# Evaluation of Candidate “Kill or Cure” Strategies to Treat MFN2-related Lipodystrophy

**DOI:** 10.1101/2025.01.01.630797

**Authors:** Ineke Luijten, Xiong Weng, Ula Kibildyte, Jana Buchan, Ami Onishi, Jake Mann, Eleanor McKay, David Savage, Robert K. Semple

**Affiliations:** Centre for Cardiovascular Science, University of Edinburgh, Edinburgh, UK; MRC Metabolic Diseases Unit, Institute of Metabolic Science, University of Cambridge, Cambridge, UK; Department of Immunology and Immunotherapy, School of Infection, Inflammation and Immunology, College of Medicine and Health, University of Birmingham, Birmingham, UK; Department of Biomedical Sciences, University of Lausanne, Lausanne, Switzerland; MRC Human Genetics Unit, Institute of Genetics and Cancer, University of Edinburgh, Edinburgh, UK

**Keywords:** Mitofusin, MFN2, lipodystrophy, multiple symmetrical lipomatosis, alcohol, rapamycin, sirolimus

## Abstract

The R707W mutation in mitofusin 2, encoded by MFN2, causes a form of Multiple Symmetrical Lipomatosis (MFN2-MSL). This resembles sporadic, alcohol-associated MSL, combining loss of lower body adipose tissue with upper body adipose hyperplasia. Morbidity and sometimes mortality arise both from mechanical complications of head and neck adipose overgrowth, and metabolic complications. We reasoned that interventions that either mitigate the underlying cellular pathology, or that exacerbate it to induce selective death of hyperplastic adipose tissue may be beneficial. We thus assessed the effect of a metabolic or pharmacologic stressors or rapamycin in *Mfn2^R707W/R707W^* mice and or derived preadipocytes. 50mmol ethanol had little effect on WT or *Mfn2^R707W/R707W^* white preadipocytes, but increased mitochondrial content and blunted mitolysosome formation in *Mfn2^R707W/R707W^*brown preadipocytes. *D*aily consumption of 20% EtOH increased brown adipose tissue mass in female *Mfn2^R707W/R707W^ mice*, and serum lactate in males. 200nM rapamycin – a candidate treatment - increased size and mitolysosome content of WT and *Mfn2^R707W/R707W^* white and brown preadipocytes, but these effects were blunted in *Mfn2^R707W/R707W^* cells. In male but not female *Mfn2^R707W/R707W^* mice, rapamycin reduced or reversed weight gain, reduced brown adipose mass, and increased serum Fgf21. Finally, a panel of other metabolic and pharmacological mitochondrial stressors solicited no selective death or ISR in *Mfn2^R707W/R707W^* preadipocytes. We conclude that ethanol mildly exacerbates MFN-MSL in mice, while rapamycin is tolerated. Lack of sensitisation to mitochondrial stressors implies that the MSL-inducing effect of MFN2 R707W may not be exerted through compromised oxidative phosphorylation.

## Introduction

We and others have described a severe and unusual form of lipodystrophy – pathological adipose tissue redistribution - caused by biallelic mutation of MFN2, encoding mitofusin 2 [1–5]. Mitofusin 2 is an outer mitochondrial membrane protein best known to play a key role in mitochondrial fusion. It also mediates tethering of mitochondria to other organelles and structures including the endoplasmic reticulum and lipid droplets [6], some of this tethering mediated by shorter MFN2 isoforms [7]. Such apposition plays an important role in processes such as apoptosis, and mitophagy.

Heterozygous MFN2 mutations most commonly cause inherited sensorimotor neuropathy [8–10]. Many causal mutations are nonsense, frameshift or essential splice site mutations, indicating that neuropathy is caused by MFN2 haploinsufficiency. MFN2-related lipodystrophy, in contrast, has only been associated with biallelic mutations, invariably including at least one R707W allele. The large majority of affected people reported have been homozygous for MFN2 R707W, although compound heterozygosity for R707W and a presumed functionally null allele is also described [1–5]. The corresponding R>W missense variant is also seen in the shorter MFN2 variants recently demonstrated to mediate endoplasmic reticulum tethering [7]. Not all patients with MFN2-related lipodystrophy have peripheral neuropathy, suggesting that penetrance and/or expressivity of MFN2 R707W-related neuropathy may be lower than that of lipodystrophy.

A fascinating and unexplained feature of MFN2-related lipodystrophy is the striking tissue- and adipose depot-selectivity of the phenotype despite ubiquitous *MFN2* expression [1–5]. As in other partial lipodystrophies, visceral adipose tissue is grossly unaffected. Subcutaneous white adipose tissue is severely affected, in contrast, with a remarkable craniocaudal pattern. Lower body, femorogluteal adipose tissue is lost, which is hypothesised but not proven to be due to loss of adipocytes. In sharp distinction, upper body white adipose tissue undergoes progressive hyperplasia and expansion [1–5]. Expansion is sometimes sufficient to occlude the airway, leading in some reported cases to premature death. This striking anatomical pattern has often led MFN2 R707W-related lipodystrophy to be called “Multiple Symmetrical Lipomatosis (MSL)” although true, encapsulated lipomas are not present.

Allied to the severe anatomical abnormality, affected people also show marked metabolic derangement, including insulin resistance, dyslipidaemia and fatty liver. Mitochondrial dysfunction is manifest as elevated serum lactate concentration, abnormal mitochondrial ultrastructure, and strong transcriptomic signatures of mitochondrial dysfunction and activation of the integrated stress response (ISR) in affected adipose tissue [1].

The combination of “mechanical” complications of upper body adipose hyperplasia and metabolic features of adipose failure imposes major morbidity. Metabolic complications can be addressed - as in other forms of lipodystrophy - by treatments inducing negative energy balance and offloading adipose tissue, but anatomical complications frequently require adipose debulking surgery. No targeted therapies are currently available, but two strategies seem worthy of consideration. Mitigating the underlying cellular pathology to slow or abrogate adipose remodelling is most appealing. One example of such an approach would be repurposing of mTOR inhibitors such as rapamycin (sirolimus), a strategy proposed for several different mitochondrial disorders based on model organism studies (e.g. [11–15]). An alternative novel approach could exploit synthetic lethality. In other words, if a treatment could be identified that selectively induces death of hyperplastic adipose tissue, then the mechanical component of the disease may be abolished. This would be at the expense of worsening deficiency of adipose tissue, but an established and growing suite of therapeutic approaches exist even for generalised adipose loss [16]. In effect, this strategy would resolve two major problems - metabolic and mechanical – into a metabolic problem alone.

To interrogate the molecular pathology of MFN2-linked lipodystrophy and facilitate preclinical translational studies we made, and have reported, *Mfn2^R707W/R707W^*mice [17]. White and brown adipose tissue (WAT and BAT respectively) from these mice recapitulated the microscopic adipose pathology described in humans, evidenced by consistent ultrastructural changes in mitochondrial morphology, transcriptomic evidence of activation of the ISR, and relative suppression of serum leptin and adiponectin. However despite prolonged challenge with high fat diet, no frank anatomical lipodystrophy was observed, and neither systemic insulin resistance nor other evidence of adipose failure was seen, attenuating the value of these mice as a disease model [17].

In this study we sought to investigate both “kill” and “cure” strategies in cells and animals to treat MFN2 R707W-related adipose dysfunction. A secondary aim was to test strategies to exacerbate the previously described phenotype of *Mfn2^R707W/R707W^* mice to enhance their utility as a disease model.

## Results

### Human dermal fibroblast studies

We have previously reported that dermal fibroblasts established from people with MFN2 R707W-related lipodystrophy do not show overt mitochondrial network disruption nor gene expression changes when cultured in standard medium containing 25 mM glucose [1]. A recent study of dermal fibroblasts from patients with sensorimotor neuropathy due to heterozygous MFN2 loss-of-function mutations also found no overt mitochondrial network abnormalities nor mitochondrial dysfunction in similar conditions [18]. When the cells were exclusively provided with galactose rather than glucose, however, mitochondrial network fragmentation and mitochondrial dysfunction were seen [18]. This was rationalised by the failure of galactose metabolism to pyruvate to generate net ATP, rendering cells dependent on mitochondrial oxidative phosphorylation when metabolising only galactose [19].

Based on these findings we assessed whether forced reliance on galactose would also unmask mitochondrial dysfunction in *MFN2^R707W/R707W^* dermal fibroblasts, thereby establishing a translationally tractable cellular disease model. We used mitochondrial morphology and transcriptional markers of the ISR as our primary readouts, based on observations of network fragmentation and a strong transcriptomic ISR signature in adipose tissue of both humans and mice homozygous for MFN2 R707W. Growth of fibroblasts from three patients in medium containing 25 mM galactose but no glucose for 48 hours did not consistently increase expression of integrated stress genes *GDF15*, *DDIT3*, or *ATF5,* and indeed GDF15 expression fell in one patient (**Figure 1A-C**). Confocal microscopy moreover revealed that although the mitochondrial network was fragmented in *MFN2^R707W/R707W^* fibroblasts, this was seen whether cultured in 25 mM glucose or 25 mM galactose-containing medium, and no difference from wild-type (WT) control cells was observed in either condition (**Figure 1D**). Galactose treatment did reduce the mitochondrial count per cell of *MFN2^R707W/R707W^* but not WT cells (**Figure 1D,E**). No differences in mean mitochondrial aspect ratio per cell were observed between genotypes or between treatments, and the mean mitochondrial branch length showed only a trend to an increase in *MFN2^R707W/R707W^* - but not WT - cells on galactose treatment (**Figure 1D,F,G**). We concluded that metabolic stress from galactose treatment was not sufficient to replicate the overt mitochondrial morphological and transcriptomic phenotype of affected adipose tissue in *MFN2^R707W/R707W^*dermal fibroblasts.

**Figure 1:**
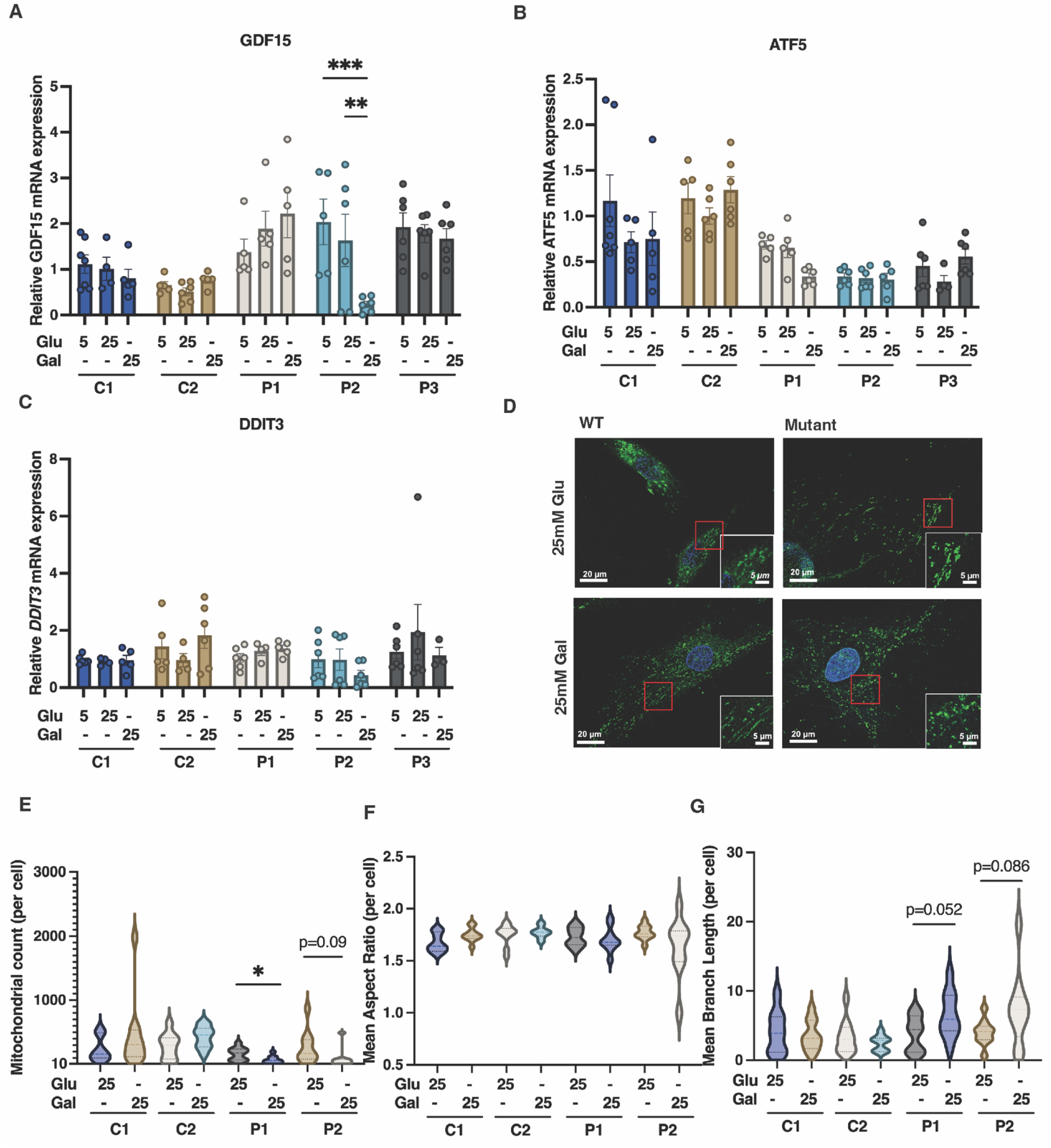
Effect of nutritional and pharmacological mitochondrial stressors on *MFN2^R707W/R707W^* and wild-type (WT) dermal fibroblasts. **A-C**: Gene expression of GDF15, DDIT3, and ATF5 in human dermal fibroblasts from MFN2 WT healthy controls (C1,2), R707W homozygous (P1,2) or R707W/delR343 compound heterozygous (P3) patients cultured in medium containing 25 mM glucose (Glu), 25 mM galactose (Gal), or 5 mM glucose for 48 hours. **D**: Representative microscopic images of mitochondria stained with MitoTracker (green) and Hoechst (blue) in human dermal fibroblasts cultured in medium containing 25 mM glucose or 25 mM galactose for 48 hours. **E-G**: Quantification of mitochondrial counts, mean aspect ratio, and mean branch length, in the same experiment. Statistical analysis was undertaken using two-way ANOVA with Tukey’s multiple comparisons test. *p<0.05, **p<0.01, ***p<0.001. N=4 for gene expression and N=10 per cell line and treatment for mitochondrial image analysis.

### Evaluation of ethanol as a mitochondrial stressor in Mfn2^R707W/R707W^ cells

We next assessed a second mitochondrial stressor, namely ethanol (EtOH), for its ability to unmask or exacerbate cellular and physiological dysfunction associated with the Mfn2 R707W mutation. The choice of EtOH was based first on the well-established deleterious effect of EtOH on mitochondrial function, mediated in part by increased redox stress and mitochondrial DNA damage [20]. A second important motivation was the observation that excess EtOH consumption is strongly associated with the sporadic form of MSL, or Madelung’s disease, that closely resembles MFN2-related lipodystrophy [21].

To enable assessment of both cellular and physiological consequences of EtOH treatment we turned to the *Mfn2^R707W/R707W^* mice [17], and murine embryonic fibroblasts (MEFs) derived from them. Initially, we assessed the effects of different concentrations of EtOH (0-100 mM) on viability of WT and homozygous MEFs. For reference, legal limits for blood alcohol when driving fall in the 10-18 mmol/L range in most countries. EtOH slightly increased the viability of homozygous MEFs, but did not affect WT cells (**Supplementary Figure 1A**). No significant genotype-nor dose-dependent differences in cell proliferation or apoptosis were observed between WT and homozygous MEFs (**Supplementary Figure 1B,C**). Determination of expression of a panel of genes related to mitochondrial biosynthesis and the ISR after 48 hours of EtOH treatment revealed only an increase in *Pgc1b* expression in *Mfn2^R707W/R707W^* cells (**Supplementary Figure 1D**), although this was present at baseline and was not exacerbated by EtOH treatment. No EtOH-induced change in expression of *Pgc1*α, nor of genes involved in mitochondrial fusion and fission (*Mfn1*, *Mfn2*, *Drp1*, *Fis1*) was seen. More tellingly, no increase in expression of ISR genes (*Ddit3*, *Trib3*, *Atf4*, *Atf5*, *Gdf15*) was seen (**Supplementary** Figure 1D), though they were sharply upregulated in adipose tissue from MFN2 R707W homozygous humans and mice [1, 17].

Our previous murine studies established that the cellular consequences of homozygosity for Mfn2 R707W are highly specific for adipose tissue, despite ubiquitous expression of Mfn2. Ultrastructural changes were discernible in preadipocytes in the earliest stages of lipid accumulation as well as in mature adipocytes [1]. MEFs do have some limited adipose differentiation capacity but are not *bona fide* preadipocytes, and so may not be a valid disease model. We thus turned next to white and brown primary preadipocytes cultured from *Mfn2^R707W/R707W^*mice. By using *Mfn2^R707W/R707W^*mice also transgenically expressing the MitoQC reporter [22, 23], we were able to assess mitochondrial network morphology in cultured cells without using exogenous mitochondrial dyes. The MitoQC construct expresses a mitochondrially-targeted tandem mCherry-GFP. The mitochondrial network thus fluoresces red and green, but upon mitophagy, when pH falls in mitolysosomes, GFP fluorescence is selectively quenched, yielding mCherry-only foci as a readout of mitophagy (**Figure 2A**).

**Figure 2:**
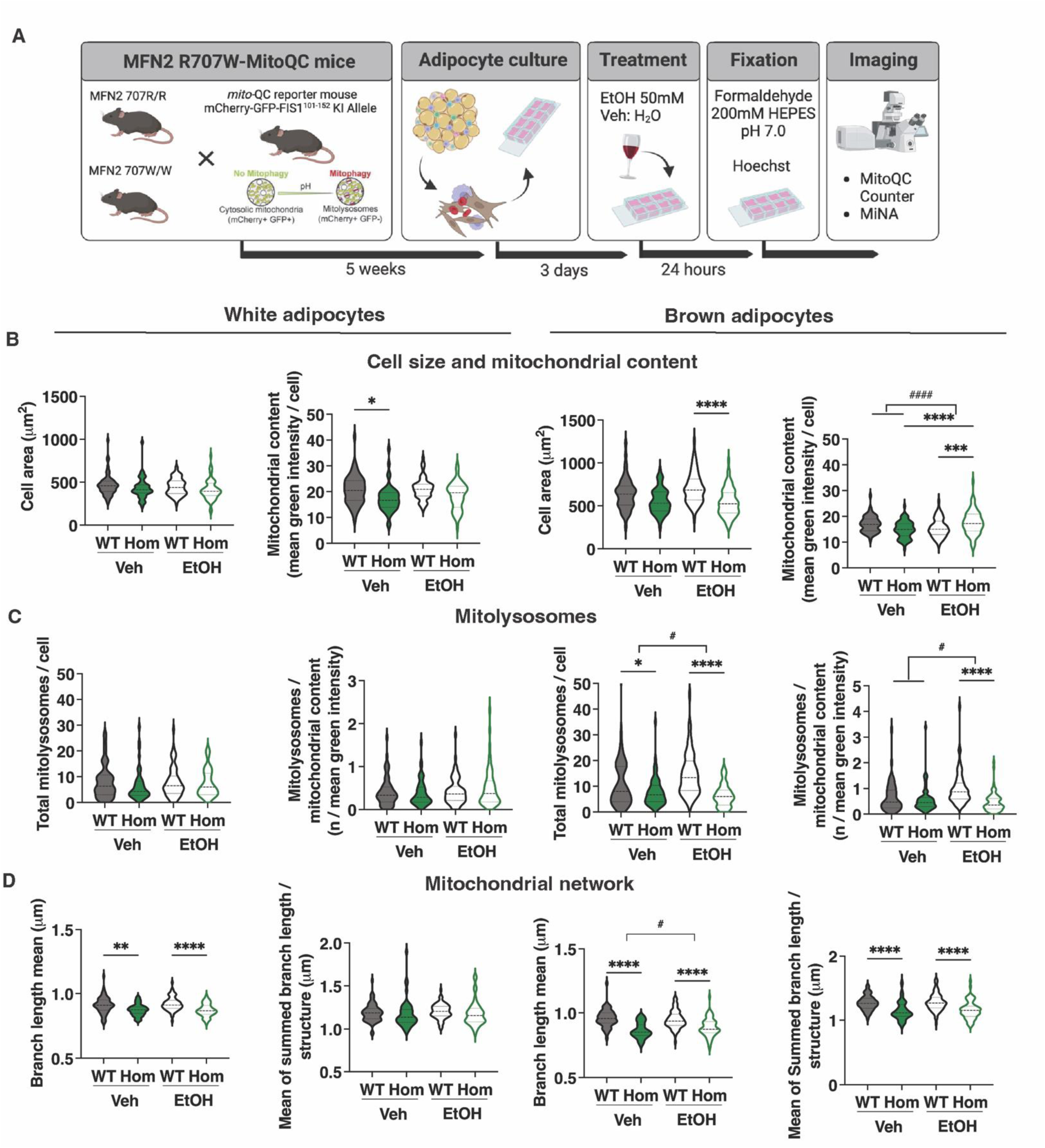
Effect of Ethanol (EtOH) on *Mfn2^R707W/R707W^* (Hom) and wild-type (WT) primary preadipocytes. **A**: Schematic overview of the Mfn2 R707W **x** mito-QC breeding strategy, primary cell culture protocol, ethanol treatment, and mitochondrial imaging analysis (Created with *BioRender*). **B**: Cell area and mitochondrial content in primary white (left panels) or brown (right panels) adipocytes isolated from WT and Hom mice, cultured for 24h in 50 mM EtOH or vehicle (Veh) **C**: Total mitolysosomes and mitolysosomes/mitochondrial content in experiment described in (B**) D**: Mean mitochondrial branch length and the mean of the summed mitochondrial branch length for each discrete mitochondrial structure in experiment described in (B). Statistical analysis was undertaken using two-way ANOVA with Tukey’s multiple comparisons test. *p < 0.05, **p < 0.01, ***p < 0.001, ****p < 0.0001; ^#^p < 0.05, ^####^ p <0.0001 for *Mfn2* genotype x treatment interaction

To test the model, we first assessed a raft of microscopic indices, including cell size, mitochondrial content, mitochondrial network morphology and mitolysosome number after exposure of primary mouse preadipocytes to carbonyl cyanide p-(trifluoromethoxy) phenylhydrazone (FCCP) (**Supplementary Figure 2A**). FCCP is a lipid-soluble protonophore that collapses the mitochondrial proton gradient, inducing disintegration of the mitochondrial network, mitophagy and ultimately cell death. In white adipocytes, 8 hours of treatment with 20μmol/L FCCP reduced mitochondrial content and mean branch length in the network, while greatly increasing the number of mitolysosomes, as expected (**Supplementary Figure 2B-E**). In brown preadipocytes, FCCP again reduced network branch length, but had equivocal or no effect on mitochondrial content and mitolysosomes per cell, which were higher at baseline than in white preadipocytes (**Supplementary Figure 2F-I**).

Having confirmed expected responses to FCCP in primary preadipocytes, we next tested the effect of 50mmol/L EtOH for 24 hours (**Figure 2A**). White preadipocytes showed only modest genotype-related changes, with reduced mitochondrial content in vehicle-treated cells only (**Figure 2B**), and reduced mitochondrial branch length in both vehicle and EtOH-treated cells (**Figure 2D**). EtOH exposure attenuated the difference in mitochondrial content seen in WT cells (**Figure 2B**), but otherwise had no discernible morphological consequences. No change in mitolysosomes was seen across genotypes and EtOH treatment (**Figure 2C**). Brown preadipocytes manifested greater morphological differences related both to genotype and to EtOH exposure. Specifically, there was a reduction in mean mitochondrial branch length and the mean sum of branch lengths in each discrete branching mitochondrial structure in *Mfn2^R707W/R707W^*cells irrespective of EtOH exposure (**Figure 2D**). Total and normalised mitolysosome content was lower in mutant than WT cells in both vehicle and EtOH-treated conditions, with the mutant appearing nearly abolishing the increases in both these indices induced by EtOH in WT cells. (**Figure 2C**). EtOH had no significant effect on mitochondrial content in WT cells, but slightly increased this in *Mfn2^R707W/R707W^*cells (**Figure 2B**). In general, morphological differences between WT and *Mfn2^R707W/R707W^* cells were exacerbated or unmasked in the presence of EtOH, with *Mfn2^R707W/R707W^* cells showing lower cell area and increased mitochondrial content (**Figure 2B**) and reduced mitolysosomes per cell (**Figure 2C**) compared to WT cells only in the presence of EtOH.

### Evaluation of the effect of ethanol in vivo

Based on evidence from mouse primary cell studies that EtOH may exacerbate some aspects of the Mfn2 R707W-related cellular phenotype, we next evaluated the effect of 2 hours per day of exposure to 20% EtOH in drinking water [24] on WT and *Mfn2^R707W/R707W^* mice over 12 weeks, starting from 12 weeks of age. WT and homozygous male mice consumed the same amount of 20% EtOH over the exposure period (**Figure 3A**), but no genotype-related differences in body weight, food intake, nor overall water intake were seen (**Figure 3B-D**). In keeping with this, no differences were seen in fat or lean mass gain (**Figure 3E,F**). BAT mass trended towards an increase in homozygous mice both at baseline and after EtOH administration, with no observed differences in inguinal (iWAT), or gonadal (gWAT) WAT, nor liver mass (**Figure 3G-J**).

**Figure 3:**
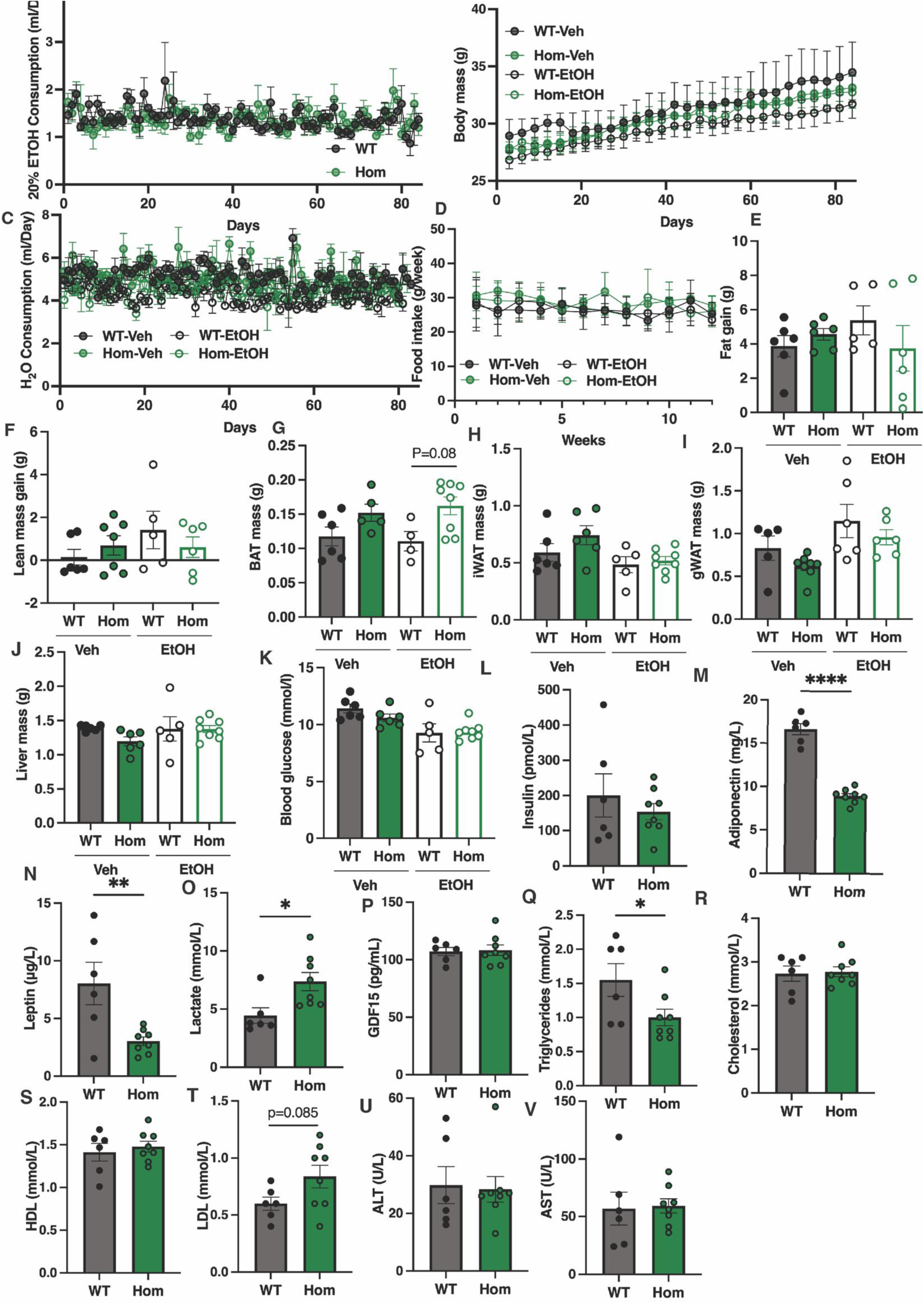
Effect of Ethanol (EtOH) on *Mfn2^R707W/R707W^* (Hom) male mice and wild-type (WT) littermates. **A-D**: 20% EtOH consumption, Body weight, water consumption, and food intake in WT and Hom males during a 3 month “Drinking in the Dark” (DID) protocol with 20% ETOH or water control. **E-J**: Analysis of fat gain, lean mass gain, brown adipose tissue (BAT), inguinal (iWAT), gonadal white adipose tissue (gWAT), and liver mass in WT and Hom males at the end of the 3 months DID protocol. **K-V**: Serum levels of glucose, insulin, adiponectin, triglycerides, leptin, lactate, total cholesterol, HDL or LDL, ALT, AST, and Gdf15 in WT and Hom males at the end of the 3 months DID protocol. Statistical analysis was performed using Two-way ANOVA with Tukey’s multiple comparisons test or Student’s t-test. *p < 0.05, **p < 0.01, ***p < 0.001. N = 4-8 per group

Serum biochemical analysis was also undertaken at the end of EtOH exposure. This showed no differences in blood glucose or plasma insulin concentrations between genotypes (**Figure 3K,L**). Serum concentrations of adipose tissue-derived adiponectin and leptin were significantly decreased in EtOH-treated homozygous mice (**Figure 3M,N**), but only to a similar extent as in untreated mice in our prior studies [17]. Serum lactate, a common index of mitochondrial dysfunction, was increased in homozygous mice, unlike prior reports of animals on chow or high fat diet alone [17], however serum Gdf15 concentration, a more general indicator of cellular stress [25], was not elevated (**Figure 3O,P**). Serum triglyceride concentration was reduced in EtOH-treated homozygous mice (**Figure 3Q**), but no changes were seen in total, HDL or LDL cholesterol (**Figure 3R,T**), nor in liver transaminases (**Figure 3U,V**).

All *in vivo* studies of EtOH exposure were undertaken also in female mice (**Supplementary Figure 3**). As in males, no divergence between WT and *Mfn2^R707W/R707W^* mice was seen in EtOH consumption, body mass, food and water consumption, and change in fat and lean mass (**Supplementary Figure 3A-F**). Also as in males, BAT mass was selectively increased (**Supplementary Figure 3G-J).** Serum biochemical studies replicated the low serum leptin and adiponectin seen in males (**Supplementary Figure 3M,N**), but lactate and triglyceride did not show any change (**Supplementary Figure 3O,Q**), unlike in males. Other analytes were also unchanged (**Supplementary Figure 3K,**L**,P,R-V**).

Collectively, our studies of the effects of exposure of *Mfn2^R707W/R707W^* preadipocytes and animals to EtOH suggest that EtOH may exacerbate some cellular and organismal manifestations of Mfn2 R707W-related (pre)adipocyte dysfunction, especially in BAT, but the effect is mild. EtOH does not lead to a frank expression of the severe biochemical and anatomical phenotype seen in humans homozygous for MFN2 R707W.

### Assessment of the effect of rapamycin on Mfn2^R707W/R707W^ cells

We next turned from environmental exposures that may worsen the MFN2 R707W lipodystrophy phenotype to assess a candidate treatment, namely the mTORC inhibitor rapamycin (sirolimus). Rapamycin is well known to upregulate mitochondrial biosynthesis, respiration, and mitophagy, thereby influencing cellular responses to nutrient and energy availability [26]. It has also been suggested as a treatment option in several primary mitochondrial disorders (e.g. [11–15]). Furthermore, footprint analysis of bulk transcriptomic data from adipose tissue of humans with *MFN2* R707W-related lipodystrophy [1] and *Mfn2^R707W/R707W^* mice [17] suggested that mTOR signalling was upregulated. Whether this was part of an adaptive response to mitigate the underlying genetic defect, or rather one of the mechanisms driving adipose hyperplasia, was not determined. This is a crucial translational question relevant to the safety of potential trials of rapamycin in people with MFN2 R707W-related lipodystrophy. Indeed, rapamycin has shown to be ineffective in some models of mitochondrial disorders [27], and to worsen clinically relevant outcomes in others [28].

Before undertaking *in vivo* studies, we first evaluated the effects of rapamycin on mitochondrial morphology and function in primary adipocytes, as before. Primary preadipocytes from WT and *Mfn2^R707W/R707W^*mice, all co-expressing MitoQC, were isolated, and treated with 200nM rapamycin or vehicle for 72 hours (**Figure 4A**). Rapamycin treatment enlarged both white and brown adipocytes, but had variable effects on mitochondrial content across genotypes and cell types, with the clearest effect being an increase in *Mfn2^R707W/R707W^*brown preadipocytes (**Figure 4B**). In keeping with its known mitophagy-inducing effect, rapamycin sharply increased total and normalised mitolysosome numbers in WT cells, with significantly attenuated increases in *Mfn2^R707W/R707W^*white and brown preadipocytes (**Figure 4C**). Effects of rapamycin on mean mitochondrial branch length and the sum of branch lengths per discrete structure were variable and small, with the only consistent effect being reduction in branch length in both white and brown WT preadipocytes (**Figure 4D**).

**Figure 4:**
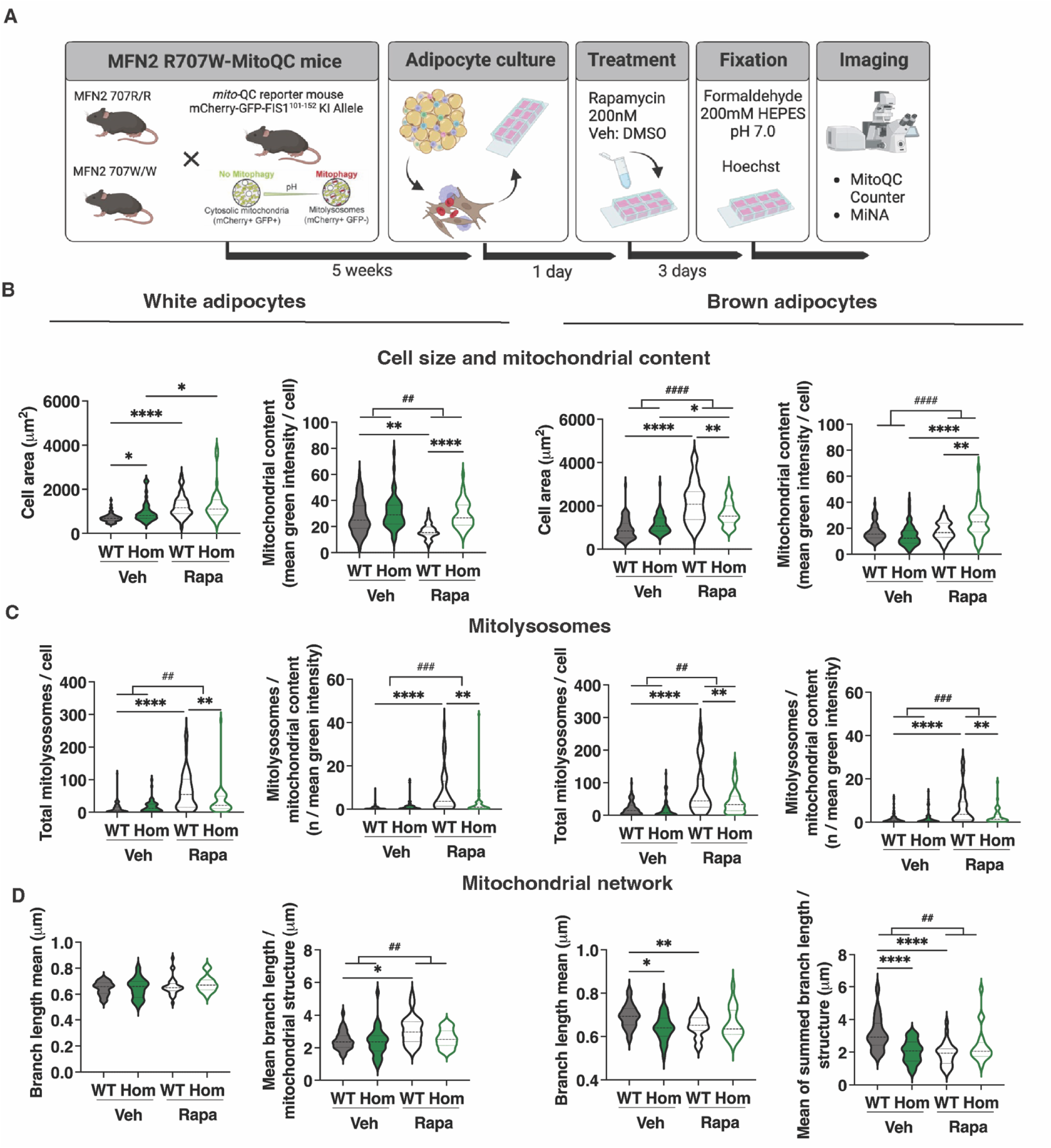
Effect of Rapamycin on *Mfn2^R707W/R707W^*and wild-type (WT) primary preadipocytes. **A**: Schematic overview of the Mfn2 R707W **x** mito-QC breeding strategy, primary cell culture protocol, rapamycin treatment, and mitochondrial imaging analysis (Created with *BioRender*). **B**: Cell area and mitochondrial content in primary white (left panels) or brown (right panels) adipocytes isolated from WT and Hom mice, cultured for 72h in 200nM rapamycin or vehicle (veh). **C**: Total mitolysosomes and mitolysosomes/mitochondrial content in experiment described in (B). **D**: Mean mitochondrial branch length and the mean of the summed mitochondrial branch length for each discrete mitochondrial structure in experiment described in (B). Statistical analysis was performed using Two-way ANOVA with Tukey’s multiple comparisons test. *p < 0.05, **p < 0.01, ***p < 0.001, ****p < 0.0001. ; ^##^p < 0.01, ^###^p < 0.001, ^####^ p <0.0001 for *Mfn2* genotype x treatment interaction

We next assessed the effects of rapamycin *in vivo*. High fat diet-fed mice were administered 8g/kg Rapamycin or vehicle intraperitoneally on alternate days for four weeks, starting three weeks into 45% high-fat diet (HFD) feeding, which in turn was started at 7 weeks. Rapamycin markedly attenuated or reversed bodyweight gain without any effect on food intake in male mice (**Figure 5A,B**). In *Mfn2^R707W/R707W^* mice, rapamycin accentuated the loss of lean mass and decrease in fat mass gain seen in WT controls, and also increased the suppression of metabolic efficiency also seen in controls (**Figure 5C,D**). No significant differences in inguinal (ingWAT) and epididymal white adipose (eWAT) mass were seen in either genotype following rapamycin treatment, but BAT mass was reduced significantly in *Mfn2^R707W/R707W^*mice only, while liver mass was unchanged (**Figure 5E,F**). Rapamycin did not change plasma insulin or blood glucose in any group, but tended to reduce serum leptin and adiponectin, although this was only significant for adiponectin in WT mice. Neither lactate nor Gdf15 was altered by rapamycin in any group, but Fgf21 was increased in *Mfn2^R707W/R707W^* mice (**Figure 5G,H**).

**Figure 5:**
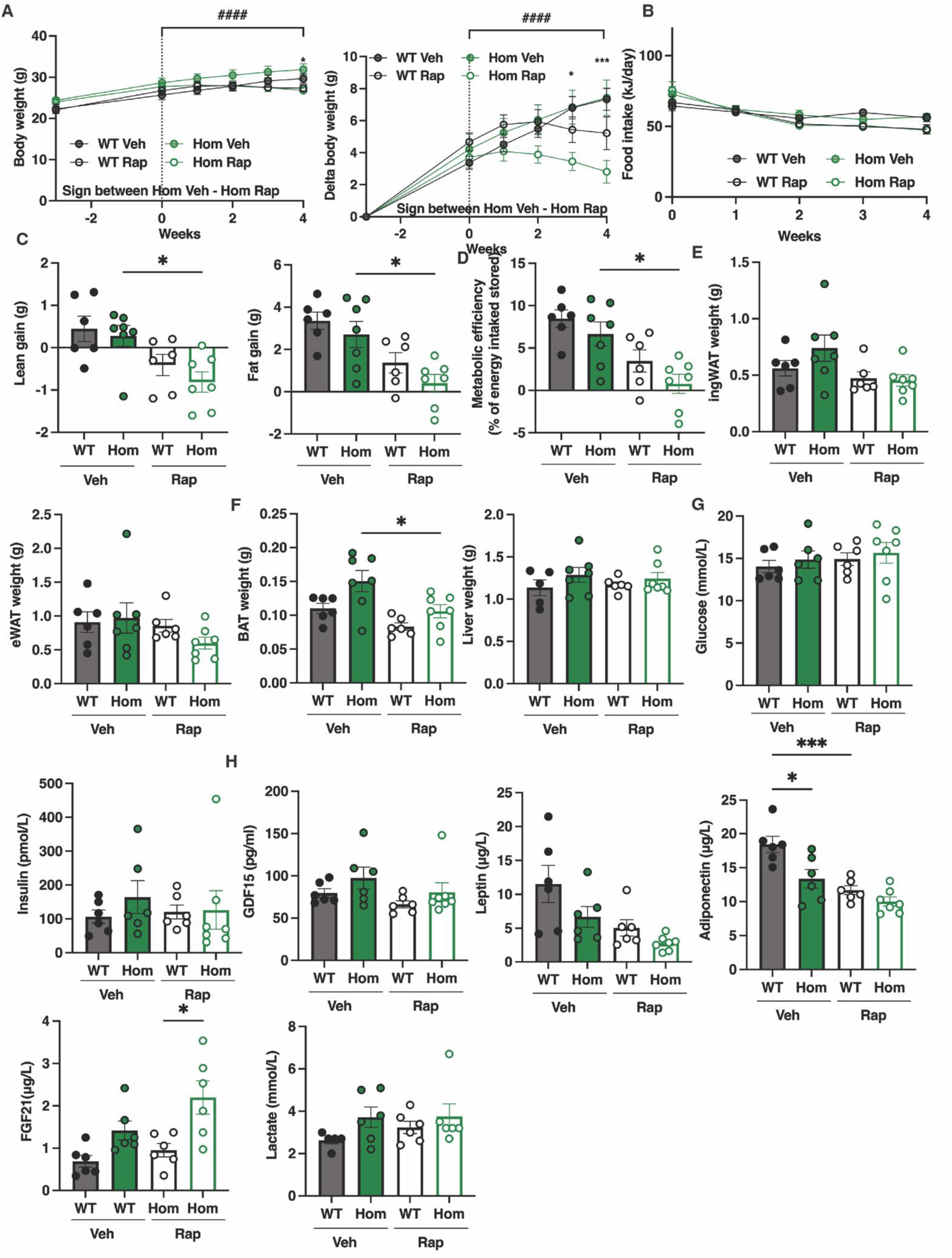
Effect of Rapamycin Treatment on male *Mfn2^R707W/R707W^*mice and wild-type (WT) littermates. **A**: Body weight and body weight gain in male mice on a 45% HFD receiving intraperitoneal injections with 8mg/kg rapamycin or vehicle (veh) every other day for 4 weeks. **B:** Food intake during rapamycin or veh treatment. **C:** Lean and fat mass gain over the 4-week treatment period. **D:** Metabolic efficiency calculated over the 4-week treatment period. **E**: ingWAT and eWAT weight at the end of the 4-week treatment period. **F**: BAT and liver weight at the end of the 4-week treatment period. **G, H**: Plasma levels of glucose, insulin, GDF15, leptin, adiponectin, FGF21, and lactate at the end of the 4-week treatment period. Statistical analysis used repeated measures two-way ANOVA with Sídák’s multiple comparisons test (A-B), and one-way ANOVA with Tukey’s multiple comparisons test (C-H). *p < 0.05, ***p < 0.001. N = 8 per group; ^####^p < 0.0001 for *Mfn2* genotype x time x treatment interaction.

The same study of rapamycin was undertaken in female mice, however almost no discernible effect of rapamycin was seen across the traits studied with the exception of a minor increase in liver weight in *Mfn2^R707W/R707W^* mice only (**Supplementary Figure 4**).

These findings do not suggest any obviously greater adverse effect on metabolism or body composition of rapamycin in *Mfn2^R707W/R707W^* mice compared to WT controls. Effects of rapamycin were more pronounced in male than female mice, and these effects appeared to be accentuated in *Mfn2^R707W/R707W^* mice.

### Assessment of further mitochondrial stressors on primary murine white preadipocytes

Having failed to observe severely deleterious or beneficial effects of EtOH and rapamycin respectively in *Mfn2^R707W/R707W^* mice, we finally undertook a pilot study to assess other strategies to exacerbate the *Mfn2^R707W/R707W^*-related cellular phenotype, both with a view to improving fidelity of the murine model to the human condition, and to establishing a proof of principle for strategies to selectively kill overgrowing adipose tissue in affected people. For this pilot study we elected to concentrate on murine white preadipocytes cultured from inguinal fat pads, given their disease relevance and the tissue selectivity of the human and mouse *Mfn2^R707W/R707W^* phenotype, using transcriptional hallmarks of the ISR as sensitive and convenient readouts of cellular pathology.

We first assessed the impact of nutritional stressors. The first of these was galactose, as in human fibroblast studies (**Figure 1**). 48 hours of growth in medium containing galactose but no glucose did not increase expression of *Atf5* and *Trib3* in cells of either genotype, and indeed *Gdf15* expression was reduced in *Mfn2^R707W/R707W^* compared to WT cells in either glucose- or galactose-containing medium, while *Atf5* and *Trib3* trended lower in *Mfn2^R707W/R707W^*cells. Only *Trib3* tended to increase in high galactose medium (**Figure 6A-D**).

**Figure 6:**
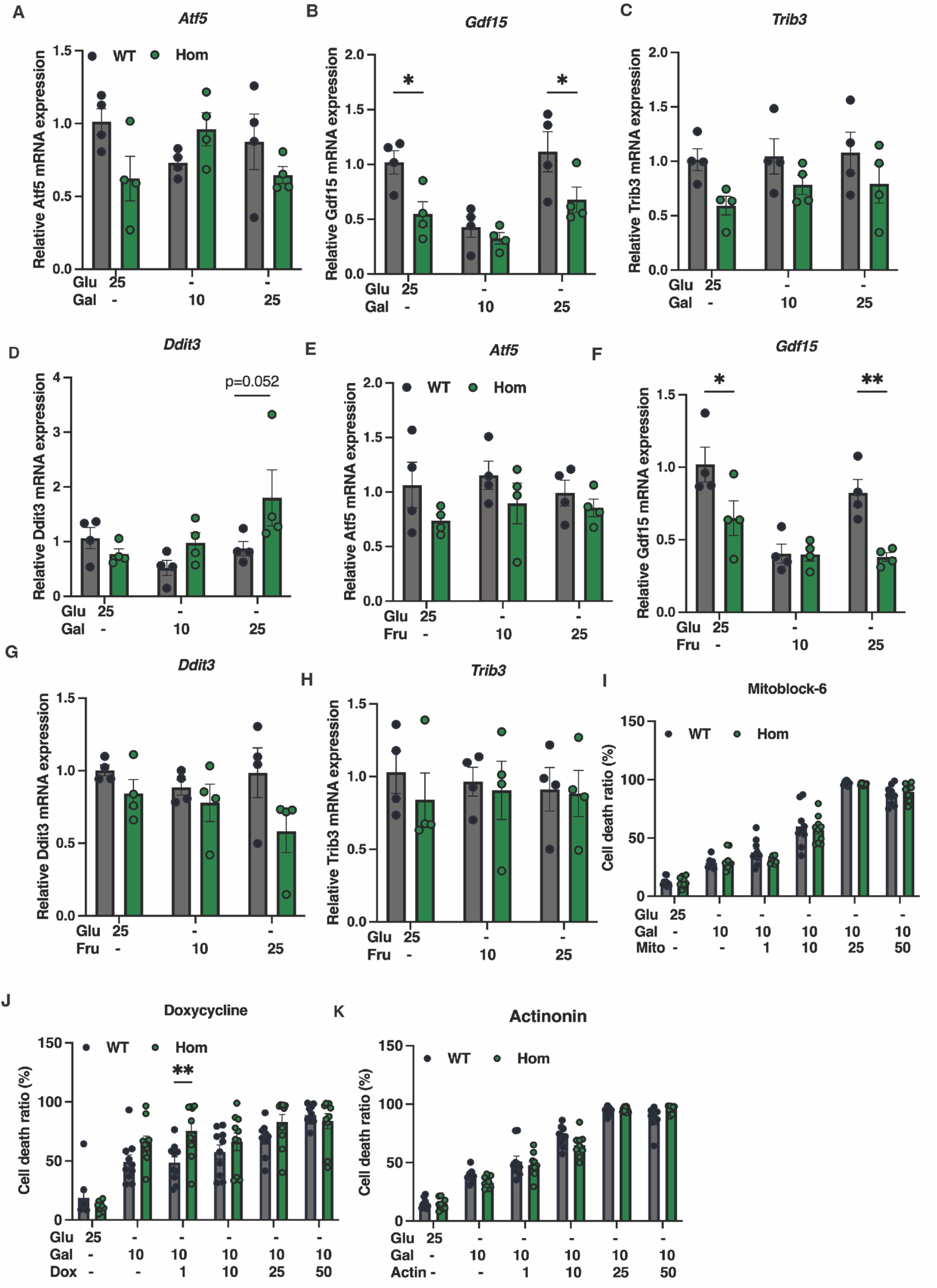
Effect of mitochondrial toxins on *Mfn2^R707W/R707W^* and wild-type (WT) primary white preadipocytes. **A-D**: Gene expression of *Atf5, Gdf15, Trib3, and Ddit3* in mouse MFN2 WT and homozygous primary white preadipocytes cultured from inguinal fat pads and treated with 10 mM, 25 mM galactose, or 25 mM glucose as a control for 48 hours. **E-H**: Gene expression of *Atf5, Gdf15, Trib3, and Ddit3* in mouse MFN2 WT and hom primary preadipocytes treated with 10 mM, 25 mM fructose (Fru), or 25 mM glucose as a control for 48 hours. **I-K**: Quantification of cell death ratios in mouse MFN2 WT and homo primary preadipocytes treated with doxycycline, Mitoblock-6, and Actinonin (N=6-10). Cells were treated with potential stressors for 48 hours in DMEM without glucose, with 1% FBS, 2 mM glutamine, and 10 mM galactose or 20 mM glucose as a control. Cells were stained with stable NucBlue® (Hoechst) for nuclei and NucGreen® Dead for dead cells at room temperature for 30 minutes. Images were taken, and the dead cell ratio was calculated as the percentage of dead cells to total cell numbers. Statistical analysis used repeated measures two-way ANOVA with Sídák’s multiple comparisons test. *p<0.05, **p<0.01. N=4 for gene expression and N=10 for cell death assay.

The second stressor tested was fructose. Forced fructose metabolism has been reported to impair mitochondrial function and activate the ISR in several tissue contexts, contributing to metabolic disturbances [29]. *SLC2A5*, encoding the GLUT5 fructose transporter, was moreover strongly upregulated in affected human adipose tissue in MFN R707W-related lipodystrophy [1]. We used a similar experimental design as for galactose, replacing glucose in culture medium with low or high concentration fructose. Fructose did not increase expression of *Ddit3*, *Atf5*, or *Trib3* in either WT or *Mfn2^R707W/R707W^* preadipocytes, and indeed, similarly to the prior experiment, *Gdf15* expression was reduced in *Mfn2^R707W/R707W^*preadipocytes exposed to either 25 mM glucose or fructose (**Figure 6E-H**).

We ended by testing three further small molecule mitochondrial stressors, using a previously reported rationale [30], and assessing them in the context of medium containing 10 mM galactose rather than glucose. In this final experiment we used cell death as the experimental readout as we were interested in potential synthetic lethality in the context of *Mfn2* R707W homozygosity. The first stressor assessed was Mitoblock-6, a small molecule that disrupts mitochondrial protein import by inhibiting Erv1/Mia40 and TIM13 [31]. Mitoblock-6 did induce dose-dependent preadipocyte death, but no Mfn2 genotype-dependency of the effect was observed (**Figure 6I**). The second stressor examined was doxycycline, a common tetracycline that inhibits mitochondrial translation, thereby provoking mitochondrial integrated stress activation in many cell types [32]. Doxycycline did not significantly increase cell death in a dose-dependent way, and no clear difference in response was discernible by genotype (**Figure 6J**). The final stressor tested was Actinonin, a small molecule that alters stability and synthesis of OXPHOS proteins [33]. Actinonin, like Mitoblock-6, provoked dose-dependent cell death, but again this effect did not show any genotype-dependency (**Figure 6K**). These findings do not support use of any of these approaches in translational strategies based on synthetic lethality.

## Discussion

MFN2 R707W, uniquely among the many pathogenic MFN2 variants described to date [34], causes major morbidity and mortality due to a combination of adipose overgrowth and metabolic complications of lipodystrophy [1–5]. There is a large unmet need to develop targeted therapy, ideally able to address both trophic and metabolic complications of the condition, but the lack of fidelity of cellular and *in vivo* disease models described to date is a barrier to translational research.

Given ubiquitous MFN2 expression, the cellular dysfunction caused by the R707W variant is remarkably selective, with only adipose tissue and to some extent peripheral nerves documented to be affected in humans [1–5]. Primary dermal fibroblasts, although expressing MFN2 strongly, appear largely normal [1], while the mouse model we previously described does not phenocopy the adipose overgrowth and systemic metabolic derangements seen in humans. It does, however, replicate key aspects of the adipose pathology, including mitochondrial network fragmentation, activation of the ISR in adipose tissue, and suppressed adipokine secretion [17].

One motivation for the current study was to assess whether additional stressors may unmask or increase cellular defects selectively in *Mfn2^R707W/R707W^* cells and mice. One of these, ethanol, did show some evidence of a genotype-selective effect, increasing brown adipose tissue mass in female *Mfn2^R707W/R707W^* mice and a trend to an increase in males, and serum lactate in male mice only. EtOH also increased or unmasked mitochondrial morphology differences between WT and mutant brown preadipocytes, suggesting that EtOH does indeed exacerbate the murine phenotype, albeit not yet offering sufficient dynamic range in the phenotype induced to serve as a robust translational disease model. This effect of EtOH is interesting given the very strong association of excess EtOH consumption with the sporadic form of MSL [21], or Madelung’s disease. The current findings suggest that deranged MFN2 function may increase susceptibility to EtOH-induced MSL.

More surprisingly, given clear evidence of mitochondrial dysfunction in adipose tissue in MFN2 R707W homozygous humans and mice, no convincing mutant-selective sensitivity was seen for either cells and animals to other, more specific, exogenous mitochondrial stressors. Stressors tested were nutritional/metabolic (forced galactose or fructose metabolism in cells) and/or pharmacological (doxycycline, mitoblock-6 or actinonin in cells). This may be an indication that impaired oxidative phosphorylation (OxPhos) is not the mechanism mediating MFN2-related lipodystrophy, even though impaired OxPhos is seen *in vivo* (evidenced by increased serum lactate, and transcriptomic signatures in affected tissue [1]). It also suggests that the relevant effect of EtOH that exacerbates the phenotype may not simply be to further impair OxPhos.

Mitochondria are often characterised simply as ATP-generating cellular “powerhouses”, but actually play a wide array of more nuanced roles. This reflects the need to adapt energy production and intermediary metabolism to different environmental conditions, and the needs of different organelles for fluxes of substrate and/or ions at different times for processes such as apoptosis and mitophagy. MFN2 subserves several functions relevant to these wider roles, mediating apposition of mitochondria to other organelles including the endoplasmic reticulum[35], enabling apoptosis and mitophagy, and lipid droplets, facilitating processes such as mitophagy and apoptosis[36]. It is plausible that the specific lipodystrophy-relevant function of MFN2 disturbed by the R707W mutation relates to tethering of mitochondria to other organelles rather than gross OxPhos dysfunction. It was recently demonstrated that two shorter splice variants of MFN2 may mediate ER tethering of mitochondria and ER morphology [7], and, interestingly, the equivalent R>W missense mutation is retained in the heptad repeat domains in each of these shorter gene products also.

The contention that it may be perturbed mitochondrial-ER interactions that underlie MFN2-related lipodystrophy remains to be tested. However the wide range of different genetic disorders of mitochondrial function now known, which feature a broad range of severity of OxPhos dysfunction [37], give further credence to the notion that there is more to MFN2 R707W-associated lipodystrophy than simply impaired OxPhos. For despite the wider range of degrees of OxPhos impairment described, lipodystrophy is reported in an extremely small subset only of mitochondrial cytopathies. These include MERRF, caused in most cases by a mitochondrial mutation in the lysyl-tRNA gene [38], and some complex disorders caused by defects in nuclear-encoded mitochondrial genes (e.g. MTX2 [39] and TOMM7 [40], both localised, like MFN2, to the mitochondrial outer membrane, and TYMP [41], a cytosolic enzyme indirectly important in maintaining mitochondrial DNA). Whether there is mechanistic commonality underpinning these select mitochondrial disorders that cause lipodystrophy is unknown but may be a fruitful line of enquiry.

The second important question addressed by this study was whether inhibition of mTOR with rapamycin is likely to be safe in MFN2 R707W-related lipodystrophy. Rapamycin had the pronounced effects on cell biology and *in vivo* metabolism expected, but we failed to see any evidence of interaction between genotype and rapamycin *in vivo* or *ex vivo*. This suggests no obvious selective toxicity of rapamycin in *Mfn2^R707W/R707W^* mice or cells, increasing confidence in the safety of any future trials.

The aims to either mitigate MFN2 R707W-relate cellular dysfunction to arrest or reverse upper body adipose overgrowth (“cure”) or to exploit synthetic lethality to selectively kill affected adipose tissue (“kill”) both remain valid. However, realising either of these strategies will require a wider, more agnostic screen of candidate interventions. As well as considering pharmacological approaches, future screens may also seek genetic modifiers of the phenotype, including knockdown of the Mfn2 paralogue Mfn1, which has been shown in some contexts to compensate for at least some neuropathy-causing mitofusin 2 mutations [42].

## Materials and methods

### Human Dermal Fibroblast Studies

Dermal fibroblasts were derived from punch skin biopsies from healthy volunteers and three patients with biallelic MFN2 mutations (P1: R707W/R343del; P2 and P3: R707W/R707W) as previously described [1]. Informed consent was obtained and confirmed in writing as part of studies approved by the UK National Research Ethics Committee (studies 18/EE/0068 and 12/EE/0405). They were cultured in DMEM (Invitrogen) supplemented with 10% fetal bovine serum (FBS) (Hyclone), 1% penicillin-streptomycin and 2 mM L-glutamine (Invitrogen), and 5 mM pyruvate, in an incubator at 37°C in 5% CO_2_/95% O_2_. For confocal studies, fibroblasts were grown on glass coverslips for 48 hours in DMEM non-glucose medium containing 25 mM glucose (Glu) or 25 mM galactose (Gal) with 1% FBS, 1% penicillin-streptomycin and 2 mM L-glutamine (Invitrogen), and 5 mM pyruvate. Cells were labeled with MitoTracker (green, Molecular Probes) and Hoechst (blue) and then fixed with 4% paraformaldehyde for 15 minutes at room temperature. Coverslips were mounted in ProLong Gold Antifade Reagent with mounting medium. Imaging and analysis were undertaken as described for MitoQC-expressing cells.

### Mouse embryonic fibroblast (MEF) culture and treatment

MEFs were derived from *Mfn2^WT/WT^*, *Mfn2^WT/R707W^* and *Mfn2^R707W/R707W^* embryos and immortalized with SV40 large T antigen as described previously [17]. All MEFs were grown at 37°C with 5% CO_2_ in Dulbecco’s Modified Eagle’s Medium (DMEM; Gibco 41966-029), containing 10% fetal bovine serum (Gibco 10500-064), Penicillin-Streptomycin-Glutamine (Gibco 10378-016), 1mM sodium pyruvate (Gibco 11360-070), MEM non-essential amino acid solution (Gibco 11140-050) and 50mM β-mercaptoethanol (Gibco 31350-010).

To assess the effect of EtOH on MEF proliferation and viability, MEFs were seeded in triplicate at 2.5*10^4^ cells/ml (c.80% confluence), grown for 24h, and then treated with the range of concentrations indicated of EtOH (absolute EtOH in H_2_O). After 24h of treatment, cell number was determined first by MTT (3-(4,5-dimethylthiazolyl-2)-2,5-diphenyltetrazolium bromide, Sigma, TOX1-1KT) assay according to the manufacturer’s instructions (Abcam ab211091). All EtOH experiments with MTT assay were repeated 6 times.

Cell number and viability after 24h EtOH treatment was also determined by Click-iT^TM^ EdU Pacific Blue^TM^ assay (ThermoFisher C10636). Growth medium was replaced with medium containing EtOH at concentrations indicated as well as 1mM EdU. After 2h incubation, cells were detached, counted and 1*10^6^ cells resuspended in 100ml Phosphate-buffered saline (PBS) containing 1% bovine-serum albumin and 400nM Apotracker^TM^ Green (Biolegend 427402). Cells were incubated for 15min at room temperature in the dark, before washing with PBS and resuspended in 100ml PBS containing 1000x diluted reconstituted LIVE/DEAD^TM^ fixable far red dead cell stain (ThermoFisher L34973). Cells were incubated for 30min at room temperature in the dark, washed, and fixed in 4% PFA. Permeabilization and Click-iT EdU detection were undertaken according to manufacturer’s instructions. Proliferation and apoptosis were determined using an Attune Nxt flow cytometer (ThermoFisher). The experiment was repeated 3 times.

### mRNA expression assays

For determination of gene expression, MEFs were washed with ice-cold PBS, harvested in TRIzol^TM^ Reagent (ThermoFisher 15596018), and RNA was extracted according to the manufacturer’s instructions. cDNA was prepared from 500ng total RNA using the High-Capacity cDNA Reverse Transcription kit (ThermoFisher 4374966). Gene expression was measured in triplicate using the primers shown (**Supplementary Table 1**) and SYBR green dye (SYBR^®^ Green JumpStart^TM^ *Taq* ReadyMix^TM^, Sigma S5193) in a LightCycler® 480 Instrument II (Roche). Specificity and efficiency of all primers was confirmed by standard dilution curve and agarose gel electrophoresis of PCR products prior to experimental reactions. Relative mRNA levels were determined using the ΔC_t_ method (2^−ΔC^) using TATA-box binding protein (Tbp) as an endogenous control.

### Animal models and husbandry

All experiments were performed under UK Home Office Project License P87539BCC, approved by the University of Edinburgh Animal Welfare and Ethical Review Board, and conducted in line with ARRIVE guidelines. Generation of mice carrying the *Mfn2* R707W mutation has been described previously [17]. *Mfn2^R707W/R707W^* mice and WT littermate controls were derived by crossing *Mfn2^WT/R707W^* mice on a C57Bl6/J background. Animals were group-housed in individually ventilated cages at 21°C with a 12h/12h light/dark cycle, controlled standard humidity (50%), and *ad libitum* access to water and chow diet (CRM, Special Diets Services). All genotyping was performed by Transnetyx (Cordova, TN).

To aid visualization of mitochondrial network structure and mitophagy, *Mfn2^WT/R707W^* mice were crossed with mice heterozygous or homozygous for the MitoQC reporter, described previously [22]. MitoQC mice were imported from MRC Harwell and housed under standard housing conditions as described above. [43]

### Mouse ethanol treatment

At 6 weeks of age, male and female *Mfn2^WT/WT^* and *Mfn2^R707W/R707W^* mice were moved to a room with reversed light/dark cycle (dark 7AM-7PM, light 7PM-7AM). After 4 weeks’ acclimatisation, mice were single caged and weekly body weight, food intake (chow diet) and water intake measurements were made. At 12 weeks of age, body composition was measured by TD-NMR (Bruker Minispec Live Mice Analyzer LF50) and mice started on a 3-month Drinking-In-the-Dark (DID) regimen [24], wherein water bottles were changed to bottles containing 20% (v/v) EtOH or vehicle (H_2_O) every day from 10-12AM. Body composition was measured monthly before the start of daily EtOH drinking. Food consumption was monitored weekly, and bodyweight every 3 days. At the end of the study, body composition was measured and a terminal bleed undertaken by cardiac puncture under general anesthesia. Tissues were harvested immediately postmortem, weighed, and snap frozen.

### Mouse rapamycin treatment

At 7 weeks of age, male and female *Mfn2^WT/WT^* and *Mfn2^R707W/R707W^* mice were single caged at 21°C and provided with *ad libitum* access to a 45% HFD (Research Diets D12541). Body weight and food intake were measured weekly throughout the experiment. At 10 weeks of age, body composition was measured by TD-NMR (Bruker Minispec Live Mice Analyzer LF50) and mice were randomly assigned to receive intraperitoneal injections with vehicle (1%DMSO, 5% PEG400, 5%Tween-80) or 8mg/kg rapamycin (VWR J62473) on alternate days for a month. After 4 weeks (at 14 weeks old), body composition was measured, and a terminal bleed undertaken by cardiac puncture under general anesthesia. Tissues were harvested immediately post-mortem, weighed, and snap frozen.

### Determination of metabolic efficiency

Food intake in kJ/day was calculated by dividing weekly food intake in g by 7 and multiplying by the following stated energy density of the diets: Chow CRM 10.74 kJ/g, HFD 19.8 kJ/g. The change in body composition during the two-week study period was calculated as (fat mass end – fat mass start) + (lean mass end – lean mass start) and converted from g to kJ using 39 kJ/g and 5 kJ/g as energy densities for fat and lean mass respectively. Metabolic efficiency was calculated by dividing the difference in body composition (in kJ) by the food intake (in kJ) over the same period, multiplied by 100.

### Serum/plasma biochemical assays

Non-fasted terminal blood samples were collected at the end of rapamycin or EtOH treatment periods. Blood glucose was measured immediately using an AccuChek Performa Nano meter. For lactate measurements, 40ul blood was collected into a fluoride EDTA tube and centrifuged immediately before freezing of plasma at –80°C. Remaining blood was allowed to clot for 15min at room temperature after which serum was isolated by centrifugation and stored at -20°C until analysis. All biochemical assays specified in **Supplementary Table 2** were undertaken by the Medical Research Council Metabolic Diseases Unit Mouse Biochemistry Laboratory, Cambridge.

### Mfn2^R707W/R707W^/MitoQC+ primary preadipocyte cultures

Primary preadipocyte cultures were made from inguinal white (ingWAT) and pooled interscapular, cervical and axillary BAT tissue depots from male *Mfn2^WT/WT^* and *Mfn2^R707W/R707W^* mice, all either homozygous or heterozygous for the MitoQC transgene. 4-6-week-old mice were sacrificed by CO_2_ and cervical dislocation, and adipose tissue depots were dissected, minced, filtered, and digested in several steps as described previously[44]. The resulting stromovascular fractions were resuspended in growth medium (DMEM (Gibco 41966-029) containing 10% newborn calf serum, 1x Penicillin-Streptomycin-Glutamine (Gibco 10378-016), 4nM insulin (Merck 19278), 10mM HEPES (Sigma H0887), 2mM glutamine (Sigma G7513), and 25mg/ml sodium ascorbate (Sigma A4034), plated on chambered coverslips (Ibidi 80806), and kept at 37°C with 5% CO_2_.

For FCCP treatment, cells were grown for 4 days during which growth medium was changed twice. On the 4^th^ day, cells were treated with FCCP (Abcam 120081) at a final concentration of 20mM or vehicle (DMSO) for 8h. For EtOH treatment, cells were grown for 3 days during which growth medium was changed once. On the 3^rd^ day, medium was replaced with medium containing 50mM EtOH or vehicle (H_2_O), and cells were incubated for 24h. For rapamycin treatment, cells were grown for 1 day after which medium was replaced with medium containing 200nM rapamycin (VWR J62473) or vehicle (DMSO). Cells were incubated for 3 days during which medium containing rapamycin or vehicle was replaced once.

### Confocal microscopy of MitoQC-expressing cells

At the end of treatment periods, primary cultures of *Mfn2^R707W/R707W^/MitoQC+* cells were rinsed twice in DPBS at 37°C and fixed in 3.7% formaldehyde in 200mM HEPES (pH7) for 15min. Cells were rinsed again and stained with 0.5mg/ml Hoechst (ThermoFisher 62249) in DPBS for 10min in the dark. After two more rinses, 4 drops of mounting medium (Ibidi 50001) were added per well, and the chambered cover slips were kept in the dark at 4°C until imaging. Cells were imaged using a Leica TCS SP8 confocal microscope with a HC PL APO 63× CS2 lens. Red and green (MitoQC), and blue (Hoechst) Z-stacks of 5-10 slices each were obtained and run through Huygens essential deconvolution software. The deconvoluted images were analyzed in Fiji using the MitoQC counter [45] and MiNA [46] macros. To analyze mitochondrial network structure, a maximum intensity z-projection of the green channel was made, and contrast was increased by 25%. Individual cells were selected and run through the MiNA macro using the default settings and the following ridge detection setup: High contrast = 0, low contrast = 40, Line width = 5, Min line length = 5. To analyze red only loci (i.e. mitophagy), a maximum intensity z-projection of the red, green and blue channel was made and brightness was increased 20% in the red and green channels. Individual cells were selected and run through the MitoQC macro using the following settings: Green channel = 2, Red channel = 1, Smoothing radius = 1, Ratio threshold = 1. Data for individual cells was compiled in Excel and around 200 cells (divided over tissues from 3 mice) were analyzed per genotype, tissue, and treatment.

### Cell death assay

To assess response to mitochondrial stressors, primary white preadipocytes were isolated from the inguinal fat pads of 6-8 weeks old *Mfn2^WT/WT^* and *Mfn2^R707W/R707W^* mice (not expressing mitoQC) and cultured as described above. To assess response to mitochondrial stressors, cells were seeded in 12 well plates at 3×10^5^cells/well. The next day medium was replaced with DMEM containing either no glucose (Thermo, 11966025), low glucose (5mM; Sigma, 50997) or galactose (10mM, 25mM; Sigma 3646739) and 1%FBS. Mitochondrial stressors (Doxycycline; Sigma 24390145, Actinonin; Sigma A6671 or mitoblock-6; Clinisciences 303215-67-0) were added to the final concentrations shown. After 48h, cell death was measured using the ReadyProbes^TM^ cell viability image kit (Blue and Green; Thermo R37609) according to manufacturer’s instruction. In brief, cells were stained by stable NucBlue® (Hoechst) for nuclei and NucGreen® Dead for dead cells at room temperature for 30mins. Images were acquired on an EVOS fluorescent microscope (Life Technologies AMF4300) and the proportion of dead cells expressed as a percentage of total cell number.

### Statistical analyses

Statistical analysis was undertaken in GraphPad Prism version 10.3.1. All tests used are indicated in figure legends, and included Student’s T test, and one way or two way ANOVA, adapted for repeated measures where appropriate. Tukey’s or Sídák’s multiple comparisons tests were used where indicated.

## Author Contributions

**IL**: conceptualization, methodology, formal analysis, investigation, writing - original draft, writing - review & editing, visualization, funding acquisition; **XW**: conceptualization, methodology, formal analysis, investigation, writing - original draft, writing - review & editing, visualization; **UK**: investigation, writing - review & editing; **JB**: investigation, writing – review & editing**; AO**: methodology, investigation, writing - review & editing. **JM**: conceptualization, resources, writing - review & editing; **EM**: investigation, writing - review & editing. **DS**: conceptualization, writing - review & editing, supervision, funding acquisition **RS**: conceptualization, resources, writing - original draft, writing - review & editing, supervision, funding acquisition.

## Supplementary Tables

**Supplementary Table 1.**
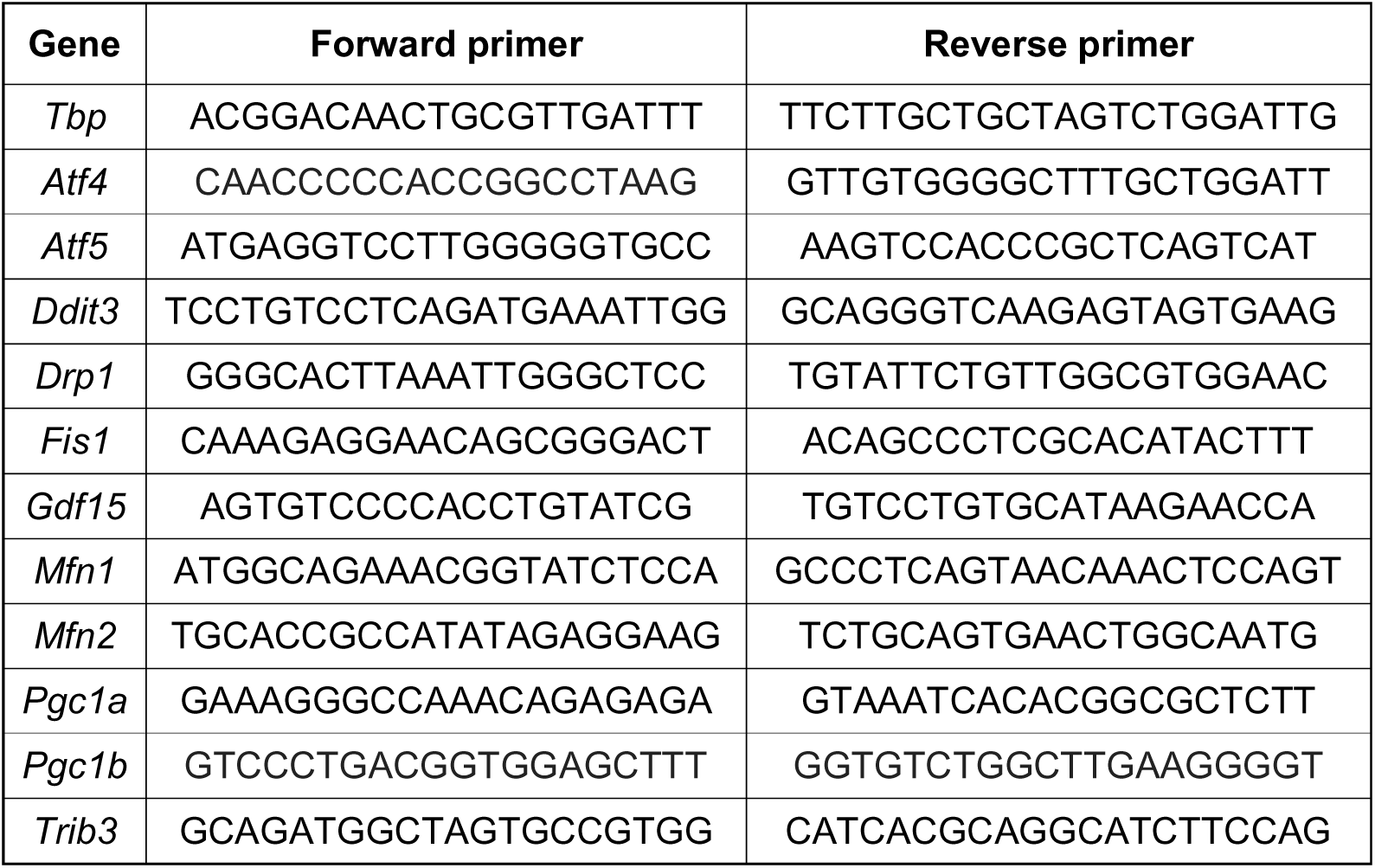
Primer sequences used for Real Time Quantitative PCR.

**Supplementary Table 2.** Biochemical assays used for *in vivo* studies.

| Analyte | Reagent manufacturer | Product code |
| --- | --- | --- |
| Insulin and leptin | MesoScale Discovery (Rockville, MD, USA) | K15124C-3 |
| Lactate | Siemens Healthcare | DF16 |
| Triglycerides | Siemens Healthcare | DF69A |
| Total cholesterol | Siemens Healthcare | DF27 |
| HDL cholesterol | Randox | CH2849 |
| LDL cholesterol | Randox | CH2656 |
| Adiponectin | MesoScale Discovery | K152BYC-2 |
| Gdf15 | R&D Systems | DuoSet ELISA (modified to run as an electrochemiluminescence assay on the Meso Scale Discovery platform) |
| Fgf21 | R&D Systems | Quantakine ELISA |
| AST | Siemens Healthcare | DF41A |
| ALT | Siemens Healthcare | DF143 |

## Supplementary Figure Legends

**Supplementary Figure 1: Effect of ethanol on wild-type and *Mfn2^R707W/R707W^*mouse embryonic fibroblasts. A:** Formazan formed in *Mfn2^WT/WT^*, *Mfn2^WT/R707W^* (707Het) and *Mfn2^R707W/R707W^* (707Hom) MEFs treated with a range of EtOH concentrations (0-100mM) for 24h expressed as a percentage of vehicle treatment. **B:** EdU positive cells in MEFs as in (A), expressed as a percentage of live cells. **C:** Apotracker positive cells in MEFs as in (A), expressed as a percentage of live cells. **D:** Gene expression for *Pgc1a, Pgc1b, Mfn1, Mfn2, Drp1, Fis1, Ddit3, Trib3, Atf4, Atf5* and *Gdf15* in MEFs as in (A), normalized to reference gene *Tbp*. Statistical analysis was performed using Two-way ANOVA with Tukey’s multiple comparisons test. *p < 0.05, **p < 0.01. N = 6, Dotted vertical lines indicate legal limits for blood alcohol when driving.

**Supplementary Figure 2: Validation of the MitoQC Reporter using FCCP. A:** Schematic overview of the Mfn2 R707W **x** mito-QC breeding strategy, primary cell culture protocol, FCCP treatment, and mitochondrial imaging analysis (Created with *BioRender*). **B,F:** Representative images of primary adipocytes isolated from ingWAT (top panels) or BAT (bottom panels) of R707W x mitoQC mice, treated with 20uM FCCP or Veh for 8h. Green: mitochondrial network, red: mitolysosomes, blue: nuclei (Hoechst). **C-E, G-I**: mitochondrial content, total mitolysosome, and branch length quantifications of the experiment described in B. Statistical analysis was performed using Two-way ANOVA with Tukey’s multiple comparisons test. *p < 0.05, **p < 0.01, ***p < 0.001, ****p < 0.0001.

**Figure 3: Effect of Ethanol (EtOH) on *Mfn2^R707W/R707W^* (Hom) female mice and wild-type (WT) littermates. A-D**: 20% EtOH consumption, Body weight, water consumption, and food intake in WT and Hom males during a 3 month “Drinking in the Dark” (DID) protocol with 20% ETOH or water control. **E-J**: Analysis of fat gain, lean mass gain, brown adipose tissue (BAT), inguinal (iWAT), gonadal white adipose tissue (gWAT), and liver mass in WT and Hom males at the end of the 3 months DID protocol. **K-V**: Serum levels of glucose, insulin, adiponectin, triglycerides, leptin, lactate, total cholesterol, HDL or LDL, ALT, AST, and Gdf15 in WT and Hom males at the end of the 3 months DID protocol. Statistical analysis was performed using Two-way ANOVA with Tukey’s multiple comparisons test or Student’s t- test. *p < 0.05, **p < 0.01, ***p < 0.001. N = 5-8 per group

**Supplementary Figure 4: Effects of rapamycin treatment on Mfn2^R707W/R707W^ female mice and wild-type littermates. A**: Body weight and body weight gain in female mice on a 45% HFD receiving intraperitoneal injections with 8mg/kg rapamycin or vehicle (veh) every other day for 4 weeks. **B**: Food intake during rapamycin or vehicle treatment. **C**: Lean and fat mass gain over the 4-week treatment period. **D:** Metabolic efficiency calculated <u>over the 4-week treatment period.</u> **E**: ingWAT and eWAT weight at the end of the 4-week treatment period. **F**: BAT and liver weight at the end of the 4-week treatment period. **G, H**: Plasma levels of glucose, insulin, Gdf15, leptin, adiponectin, FGF21, and lactate at the end of the 4- week treatment period. Statistical analysis was performed using repeated measures two-way ANOVA with Sídák’s multiple comparisons test (A-B), and one-way ANOVA with Tukey’s multiple comparisons test (C-H). **p < 0.01. N = 8 per group.

