## Supplementary Figures for "Evaluation of Candidate “Kill or Cure” Strategies to Treat MFN2-related Lipodystrophy"

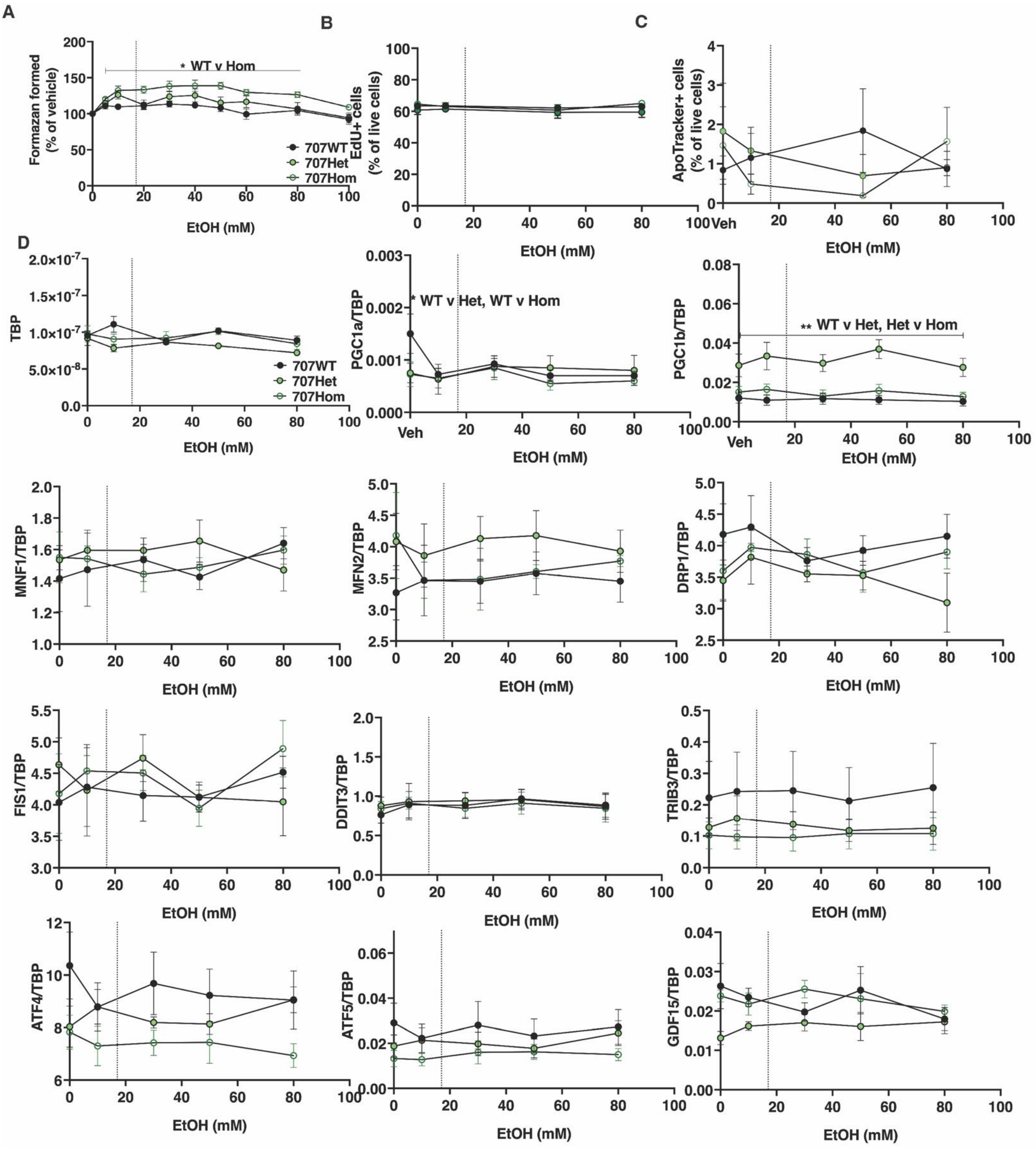


**Supplementary Figure 1. Effect of ethanol on on wild-type and *Mfn2*$^{R707W/R707W}$ mouse embryonic fibroblasts**

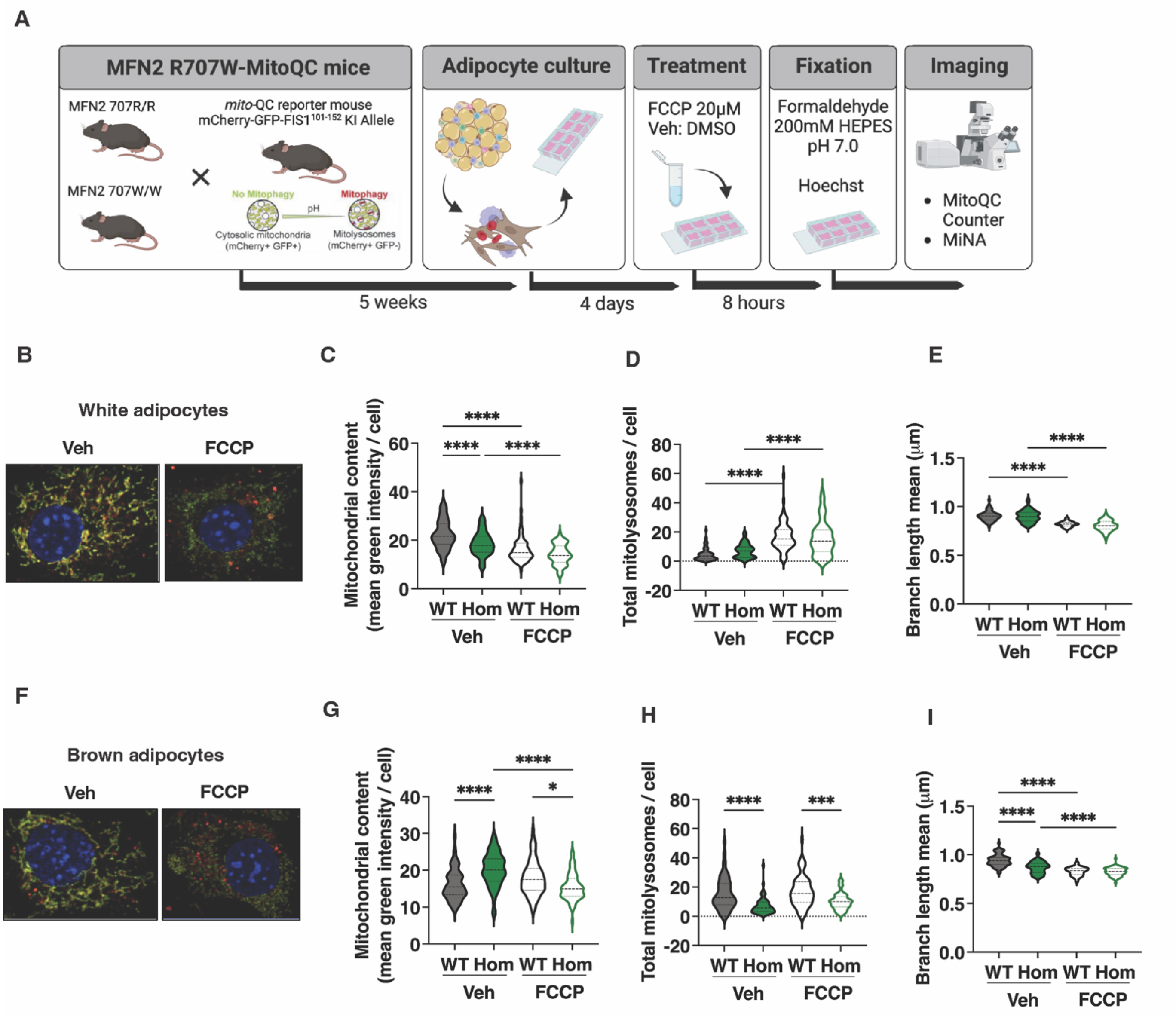


**Supplementary Figure 2: Testing of MitoQC Reporter using FCCP.**

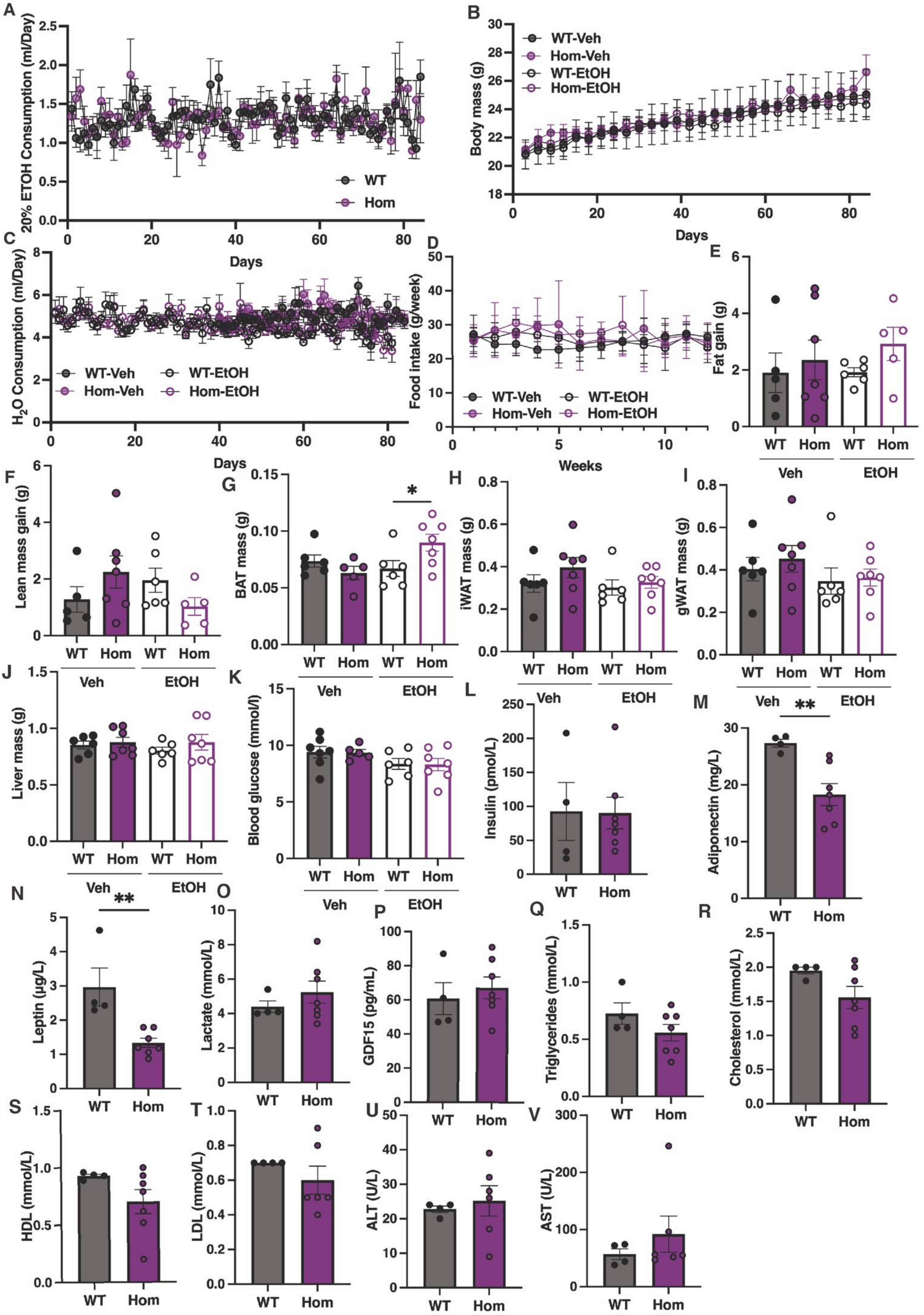


**Supplementary Figure 3. Effect of ethanol on female wild-type and *Mfn2*$^{R707W/R707W}$ mice**

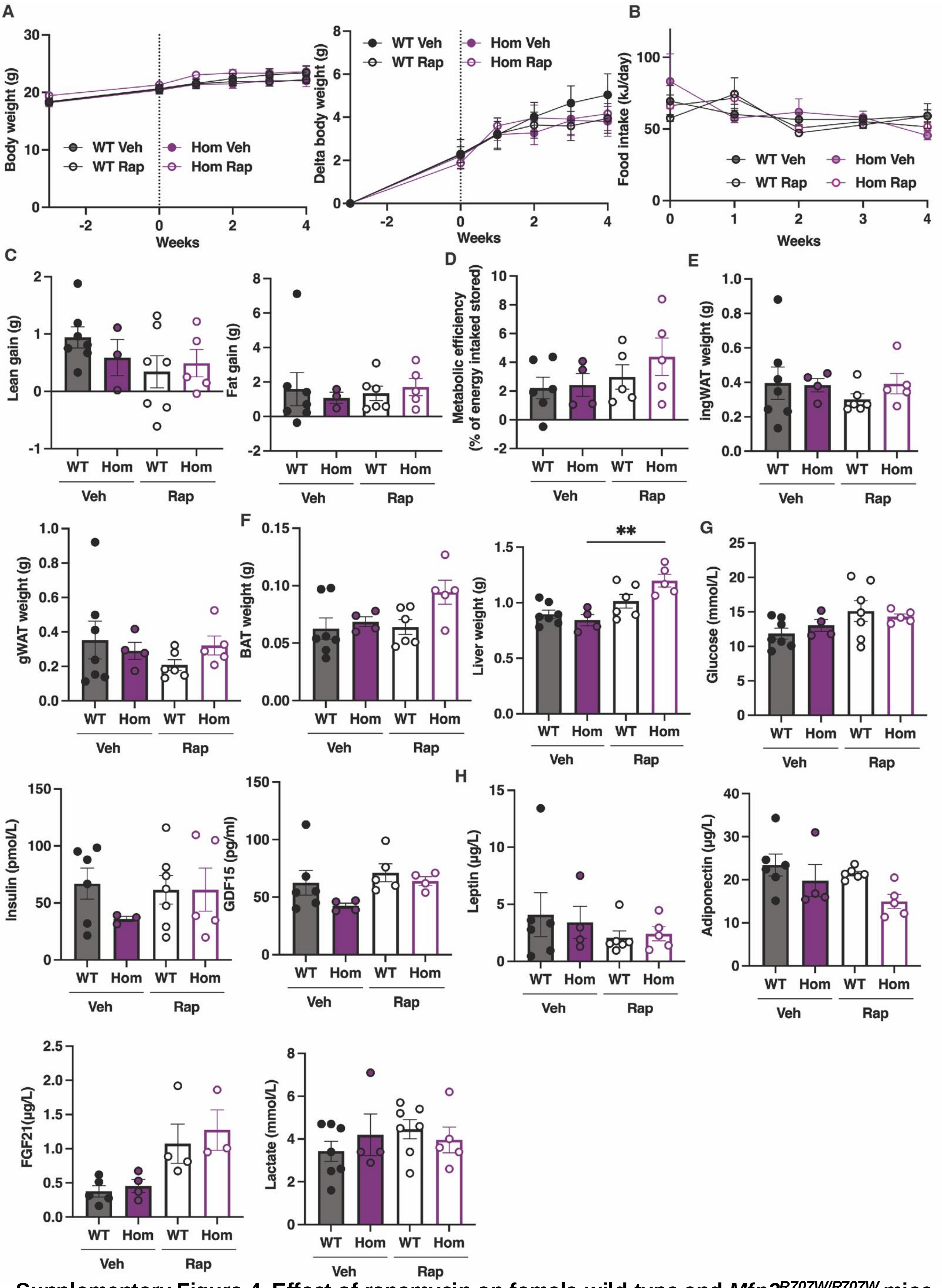


**Supplementary Figure 4. Effect of rapamycin on female wild-type and *Mfn2$^{R707W/R707W}$* mice**
